# The mechanism underlying the organization of Borna disease virus inclusion bodies is unique among mononegaviruses

**DOI:** 10.1101/2021.05.24.445377

**Authors:** Yuya Hirai, Keizo Tomonaga, Masayuki Horie

## Abstract

Inclusion bodies (IBs) are characteristic biomolecular condensates organized by mononegaviruses. Here, we characterize the IBs of Borna disease virus 1 (BoDV-1), a unique mononegavirus that forms IBs in the nucleus, in terms of liquid-liquid phase separation (LLPS). The BoDV-1 phosphoprotein (P) alone induces LLPS and the nucleoprotein (N) is incorporated into the P droplet *in vitro*. In contrast, co-expression of N and P is required for the formation of IB-like structure in cells. Furthermore, while BoDV-1 P binds to RNA, an excess amount of RNA dissolves the liquid droplets formed by N and P. Notably, the N-terminal intrinsically disordered region of BoDV-1 P is essential to drive LLPS and bind to RNA, suggesting that both abilities could compete with one another. These features are unique among mononegaviruses, and thus this study will contribute to a deeper understanding of LLPS-driven organization and RNA-mediated regulation of biomolecular condensates.

## Introduction

The order *Mononegavirales* contains many pathogens of great importance to public health, such as measles virus, Ebola virus, rabies virus, and human respiratory syncytial virus (RSV) ^1^. These mononegaviruses form various types of inclusion bodies (IBs) in infected cells, which are thought to be the sites of viral replication ^2^. The IBs of mononegaviruses are membraneless organelles, also called biomolecular condensates ^3–6^. Growing evidence has shown that the IBs of several mononegaviruses, such as rabies virus ^7^, vesicular stomatitis virus (VSV) ^8^, measles virus ^9,10^, and RSV ^11^, are organized by liquid-liquid phase separation (LLPS).

The viral ribonucleoprotein (vRNP) complexes of mononegaviruses, which function as the fundamental units of transcription and replication, consist of viral genomic RNA, nucleoprotein (N), phosphoprotein (P), and large RNA-dependent RNA polymerase (L). Among the components of vRNP complexes, the expression of N and P is sufficient for the formation of IBs of rabies virus ^7^, measles virus ^9^, RSV ^11^, human metapneumovirus ^12^, and human parainfluenza virus 3 ^13^. Previous studies of these IBs have provided insights into the viral replication strategies, thereby contributing to the control of viral infectious diseases. Additionally, since viruses exploit the host machinery for replication, further studies would be helpful to gain a deeper understanding of the molecular condensates in cells.

Borna disease virus 1 (BoDV-1) is a prototype virus of the family *Bornaviridae* of the order *Mononegavirales*, which causes fatal encephalitis in mammals, including humans ^14^. BoDV-1 is a unique mononegavirus as replication occurs in the nucleus of the infected cell, whereas that of most mononegaviruses occurs in the cytoplasm ^15^. Consequently, the IBs of cytoplasmic mononegaviruses are formed in the cytoplasm, while those of BoDV-1, termed as viral speckle of transcripts (vSPOTs), are organized in the nucleus ^16^. The vSPOTs are membraneless spherical structures containing the viral genome, antigenome, and several viral proteins. The BoDV-1 genome encodes six genes, N, P, matrix protein (M), envelope glycoprotein (G), L, and accessory protein (X), among which N, P, M, L, and X are localized in the vSPOTs ^17,18^.

Although present in the nucleus, the vSPOTs share common properties with those of the other mononegaviruses, such as membraneless structures, and their components. However, the surrounding environment is important for the formation of biomolecular condensates. As described above, BoDV-1 forms IBs in the cell nucleus, where the surrounding environment differs from that of the cytoplasm. Thus, the IBs of BoDV-1 may be formed by a different mechanism from those of other cytoplasmic mononegaviruses. Nonetheless, the propensities and minimal components of BoDV-1 IBs remain entirely elusive.

In this study, to clarify the mechanism of the formation of vSPOTs, we analyzed the properties of the BoDV-1 N and P proteins from the aspect of phase-separated condensates. We revealed that the BoDV-1 P protein alone causes LLPS *in vitro*, depending on the intrinsically disordered region (IDR) present at the N-terminal region. Although BoDV-1 N alone is not sufficient for the induction of LLPS, it was incorporated into the BoDV-1 P droplets. Also, co-expression of BoDV-1 N and P is sufficient to induce the formation of IB-like structures in cells. Further, P of BoDV-1, but not VSV or RSV, binds to RNA. Interestingly, the N-terminal IDR of BoDV-1 P is involved in this RNA-biding activity, which is overlapped with the region important for driving LLPS, while an excess amount of RNA dissolved the liquid droplets consisting of BoDV-1 P alone and BoDV-1 N and P. These findings are expected to help clarify the fundamental mechanism of the morphological regulation of the IBs of BoDV-1 and the comprehensive mechanisms underlying the formation of biomolecular condensates.

## Materials and methods

### DNA constructs and antibodies

*Escherichia coli* expression vectors were constructed as follows. Oligonucleotides of codon-optimized BoDV-1 N, P, His-tagged enhanced green fluorescent protein (EGFP), and His-tagged mCherry (strain He/80/FR) were synthesized by Thermo Fisher Scientific (Waltham, MA, USA). The synthesized oligonucleotides of BoDV-1 N and P were inserted into the *Xho*I and *Bam*HI restriction sites of the plasmid pET-15b using NEBuilder HiFi DNA Assembly Master Mix (E2621; New England Bioscience, Ipswich, MA, USA), which were designated as pET-15b-BoDV-1-Nco and pET-15b-BoDV-1-Pco, respectively. The synthesized oligonucleotides of His-tagged mCherry and EGFP were cloned into pET-15b using the *Nco*I and *Nde*I restriction sites, which were named pET-15b-His-mCherry and pET-15b-His-EGFP, respectively. The codon-optimized N and P sequences were amplified by PCR and then inserted into pET-15b-His-mCherry and pET-15b-His-EGFP, respectively, using the *Nco*I site.

For mammalian expression vectors, the coding regions of the N and P genes of BoDV-1 (strain He/80/FR) with the Kozak sequence were amplified by PCR, and then cloned into the pCAGGS vector at the *Eco*RI and *Xho*I restriction sites using NEBuilder HiFi DNA Assembly Master Mix. All of the primer and plasmid sequences are available upon request.

The following antibodies were used in this study: anti-BoDV-1 N mouse monoclonal antibody (HN132), anti-BoDV-1 P rabbit polyclonal antibody (HB03), anti-high mobility group box 1 (HMGB1) antibody (sc-74085; Santa Cruz Biotechnology, Inc., Dallas, TX, USA), goat anti-mouse immunoglobulin G (IgG) (H+L) highly cross-adsorbed secondary antibody, Alexa Fluor 488 (A-11029; Thermo Fisher Scientific), goat anti-mouse IgG (H+L) highly cross-adsorbed secondary antibody, Alexa Fluor 568 (A-11031; Thermo Fisher Scientific), goat anti-rabbit IgG (H+L) highly cross-adsorbed secondary antibody, Alexa Fluor 488 (A-11034; Thermo Fisher Scientific), and goat anti-rabbit IgG (H+L) highly cross-adsorbed secondary antibody, Alexa Fluor 568 (A-11036; Thermo Fisher Scientific).

### Protein expression and purification

N-terminal His-tagged recombinant proteins were expressed in Rosetta™ 2(DE3) pLysS competent cells (71403; Novagen, Inc., Madison, WI, USA) by induction with 0.5 mM isopropyl β-d-1-thiogalactopyranoside at 30°C for 4 h (N, P, and P’), at 25°C for 5 h (N-mCherry, VSV P, and RSV P), or at 16°C overnight (P-EGFP). The pelleted cells were resuspended in lysis buffer (20 mM Tris-HCl, pH 7.4, 500 mM NaCl, 10 mM imidazole) containing a protease inhibitor cocktail (25955-24; Nacalai Tesque, Inc., Kyoto, Japan). After sonication, the lysates were clarified by centrifugation at 20,000 × *g* for 10 min at 4°C. Then, the supernatants were loaded onto a column containing TALON® Metal Affinity Resin (635501; Takara Bio, Inc., Shiga, Japan). The resins were washed with lysis buffer and the His-tagged proteins were eluted with elution buffer (20 mM Tris-HCl, pH 7.4, 500 mM NaCl, 300 mM imidazole), dialyzed with dialysis buffer (20 mM Tris-HCl, pH 7.4, 500 mM NaCl), and concentrated with Amicon® Ultra Centrifugal Filters (Merck KGaA, Darmstadt, Germany).

### *In vitro* phase separation assay

The concentrated proteins were diluted to the indicated concentrations in phase separation buffer (20 mM Tris-HCl, pH 7.4, 150 mM NaCl, except for Fig. 1D). RNA and 1,6-hexanediol (H0099; Tokyo Chemical Industry, Tokyo, Japan) were added to the mixed reaction samples at final concentrations of 200 ng/µl and 6%, respectively. Fluorescently labeled liquid droplets were formed by mixing 9.5 µM P and 0.5 µM P-EGFP (P droplets), and 5 µM N, 4.5 µM P, and 0.5 µM P-EGFP (NP droplets). The mixed samples were transferred to the wells of 384-well EZVIEW® glass-bottom assay plates (5883-384; Iwaki Co., Ltd., Chiba, Japan) and observed at room temperature (RT) with a laser scanning confocal microscope (LSM 700; Carl Zeiss AG, Oberkochen, Germany) equipped with a Plan-Apochromat 63X objective lens (numerical aperture = 1.4). The size and intensity of each droplet was calculated using ImageJ software (https://imagej.nih.gov/ij/). The sizes and intensities are presented as the average of at least three separate fields.

**Fig. 1.**
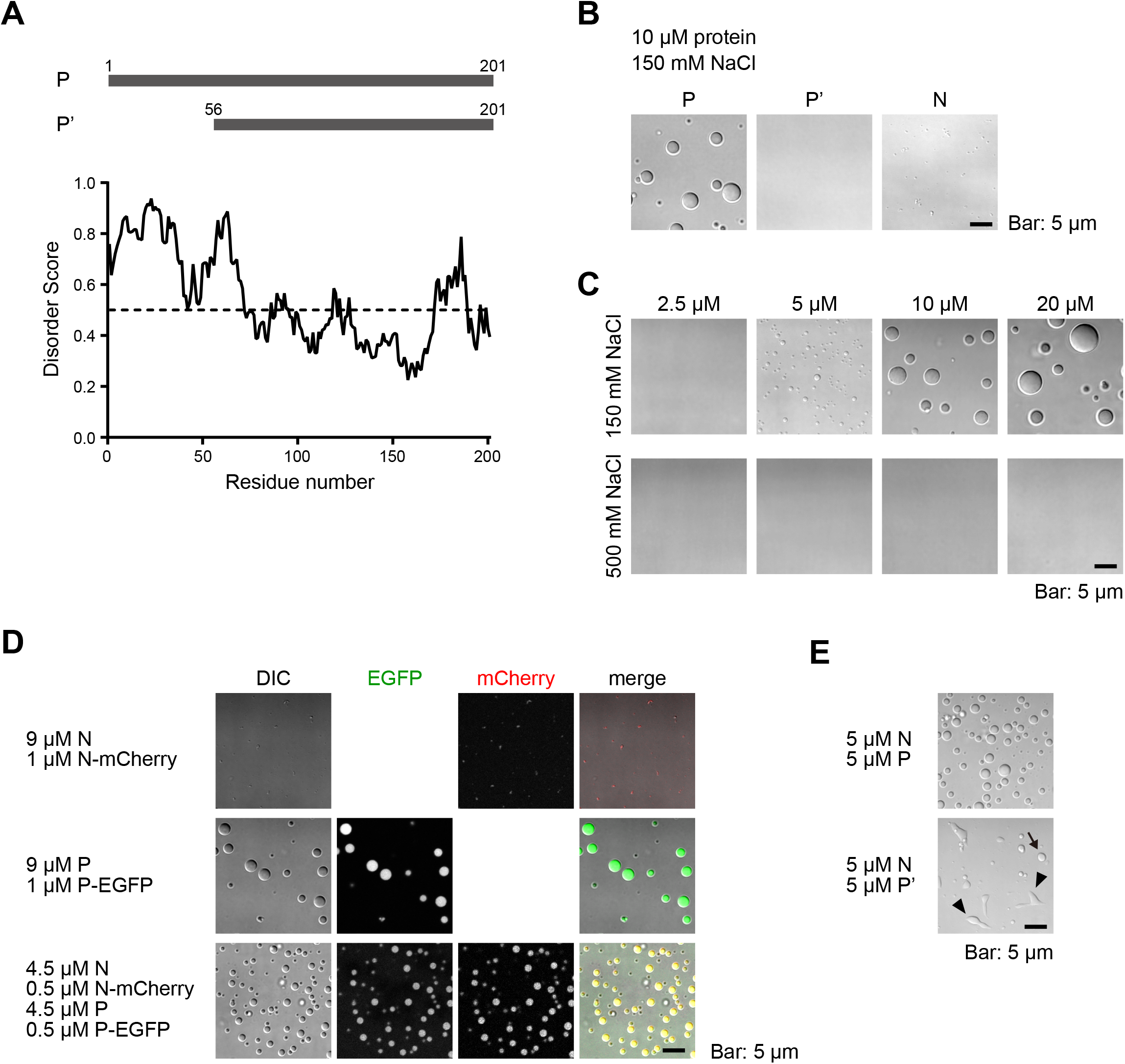
IDR-dependent formation of liquid droplets by BoDV-1 N and P. **A**. A schematic diagram of the amino acid regions of p24 and p16. The IDRs were predicted by IUPred2A ^39^. **B**. Differential interference contrast (DIC) images of the liquid droplet formation of BoDV-1 P (p24), P (p16), and N. **C**. DIC images of the liquid droplet formation of the indicated protein concentration of BoDV-1 P (p24) in 150 mM and 500 mM NaCl. **D**. DIC and fluorescence images of the liquid droplet formation of the indicated protein combination. **E**. DIC images of the liquid droplet formation of the indicated protein combination. The arrowheads indicate the distorted aggregate-like structures and the arrow indicate the droplets that were scarcely observed.

### *In vitro* transcription

DNA templates (a BoDV-1 minigenome containing the complementary sequence of Gaussia luciferase [Gluc] flanked by the leader and trailer sequences of the BoDV-1 genome [Fig. S3A] or the complementary sequence of Gluc) for *in vitro* transcription were amplified by PCR using primers containing the T7 promoter sequence at the 5’-end and pBDVmg-Gluc^19^ as a template. The *in vitro* transcription reaction was performed using T7 RNA polymerase (2540A; Takara Bio, Inc.) in accordance with the manufacturer’s protocol. To label RNA, Cy3-UTP (ENZ-42505; Enzo Life Sciences, Inc., Farmingdale, NY, USA) was added into the *in vitro* transcription reaction at a final concentration of 0.05 mM. The transcribed RNA was treated with DNaseI, then purified using the phenol/chloroform isolation method and resuspended in RNase-free water.

### Electrophoretic mobility shift assay

Proteins at the indicated concentrations were mixed with 2 µg of RNA, incubated for 30 min at RT, and then separated by 1.5% agarose gel electrophoresis. RNA signals were detected using GelGreen®, a highly sensitive, non-toxic green fluorescent nucleic acid dye (41004; Biotium, Inc., Fremont, CA, USA).

### Cell culture

Human oligodendroglioma (OL) cells persistently infected with BoDV-1 strain huP2br ^20^ were cultured in high-glucose (4.5%) Dulbecco’s modified Eagle’s medium (DMEM) (11965092; Thermo Fisher Scientific) supplemented with 5% fetal bovine serum. U-2 osteosarcoma (OS) cells (92022711; European Collection of Authenticated Cell Cultures, Public Health England, London, England) were cultured in DMEM (08456; Nacalai Tesque, Inc.) supplemented with 10% fetal bovine serum. The cells were transfected using Lipofectamine 2000® Transfection Reagent (11668027; Thermo Fisher Scientific) or Polyethylenimine “Max” (24765-1; Polysciences, Inc., Warrington, PA, USA).

### Immunofluorescence microscopy

The cells were cultured on coverslips, fixed with 4% paraformaldehyde for 10 min, blocked with 5% bovine serum albumin containing 0.5% Triton X-100 for 15 min, and then probed with primary antibodies for 2 h at RT. Afterward, the cells were washed twice with phosphate-buffered saline (PBS), incubated with the secondary antibodies and 4′,6′-diamidino-2-phenylindole for 1 h at RT, then washed three times with PBS, mounted with ProLong® Diamond Antifade Reagent (Life Technologies, P36961), and observed using a laser scanning confocal microscope (LSM 700; Carl Zeiss AG) equipped with a Plan-Apochromat 63× objective lens (numerical aperture = 1.4).

### Time-lapse imaging of BoDV-1-infected cells

BoDV-1-infected OL cells transfected with a plasmid expressing EGFP-tagged P ^21^ were cultured in a glass-bottom dish and imaged with a C1 confocal laser scanning microscope (Nikon Corporation, Tokyo, Japan) equipped with a CFI Apo Lambda S 60× objective lens (numerical aperture = 1.4), a 37°C heating stage, and lens heater. A series of images were acquired at intervals of 10 s.

### Fluorescence recovery after photobleaching (FRAP) analysis

To induce LLPS, 9 µM N, 1 µM N-mCherry, 9 µM P, and 1 µM P-EGFP were mixed. FRAP analysis was performed using a laser scanning confocal microscope (LSM 700; Carl Zeiss AG). After acquiring 10 images (97.75 ms per frame), the indicated spots were bleached using a 488-nm laser with 15 iterations. Then, 590 images were captured. The fluorescence intensity was quantified using ZEN software (Carl Zeiss AG).

## Results

### P triggers and is necessary for the formation of liquid droplets of BoDV-1 N and P

We first determined the viral protein(s) that forms liquid droplets *in vitro*. As described above, the N and (partial) P proteins are necessary for the formation of liquid droplets by other mononegaviruses ^10,11^. In addition, the IDR is one of several factors that initiate phase separation ^4,22,23^. Therefore, we predicted the IDRs of the BoDV-1 N and P proteins (Fig. 1A and S1). P was predicted to have a disordered amino acid sequence, especially at the N-terminal region (Fig. 1A). Notably, the BoDV-1 genome encodes two P isoforms, P and P’, which are translated from alternative start codons ^24^. The fully disordered region was predicted to be present only in P, but not P’ (Fig. 1A).

To investigate which proteins cause phase separation, we examined whether P, P’, and/or N form liquid droplets *in vitro*. At a near physiological salt concentration (150 mM NaCl), P clearly formed liquid droplets at a protein concentration of 10 µM (Fig. 1B), while N formed aggregate-like structures but not liquid droplets, and P’ formed neither (Fig. 1B). At a protein concentration of 5 µM, but not 2.5 µM, in 150 mM NaCl, P formed liquid droplets (Fig. 1C). However, in 500 mM NaCl, P failed to initiate a phase separation, even at a protein concentration of 20 µM (Fig. 1C).

In infected cells, N and P are colocalized within viral biomolecular condensates, vSPOTs ^16^. Therefore, we next investigated the effects of P on the aggregate-like structures of N by mixing both N and P *in vitro*. Here we mixed fluorescently labeled proteins (N-mCherry and P-EGFP, both were tagged at the C-terminus) to the reaction mixture to visually distinguish between the N and P proteins. The fluorescent protein-labeled N and P were incorporated into droplets formed by unlabeled N and P (Fig. 1D), respectively, indicating that both fluorescent protein-labeled N and P exhibited the same behaviors as the unlabeled proteins. When N and P were mixed, the liquid droplets were observed, which include not only P but also N (Fig. 1D). However, a mixture of N and P’ formed distorted aggregate-like structures with few liquid droplets. (Fig. 1E).

These results suggest that P, but not P’, alone can form liquid droplets via the IDR at the N-terminal region and N forms liquid droplets with P, which is also dependent on the IDR of P.

### Viral condensates formed by BoDV-1 have liquid-like properties

BoDV-1 forms membraneless IBs containing BoDV-1 N and P in the nucleus of the infected cells. Also, we showed that BoDV-1 N and P formed liquid droplets *in vitro* (Fig. 1D and E). These properties are similar to those of cellular biomolecular condensates. However, the detailed characteristics of BoDV-1 IBs remain unclear. Therefore, we examined the potential liquid properties of BoDV-1 IBs. Time-lapse imaging of BoDV-1-infected cells revealed the fusion of two vSPOTs (Fig. 2A). Considering BoDV-1 P is continuously exchanged between vSPOTs ^25^, this result indicates that vSPOTs, like other biomolecular condensates, also have liquid properties.

**Fig. 2.**
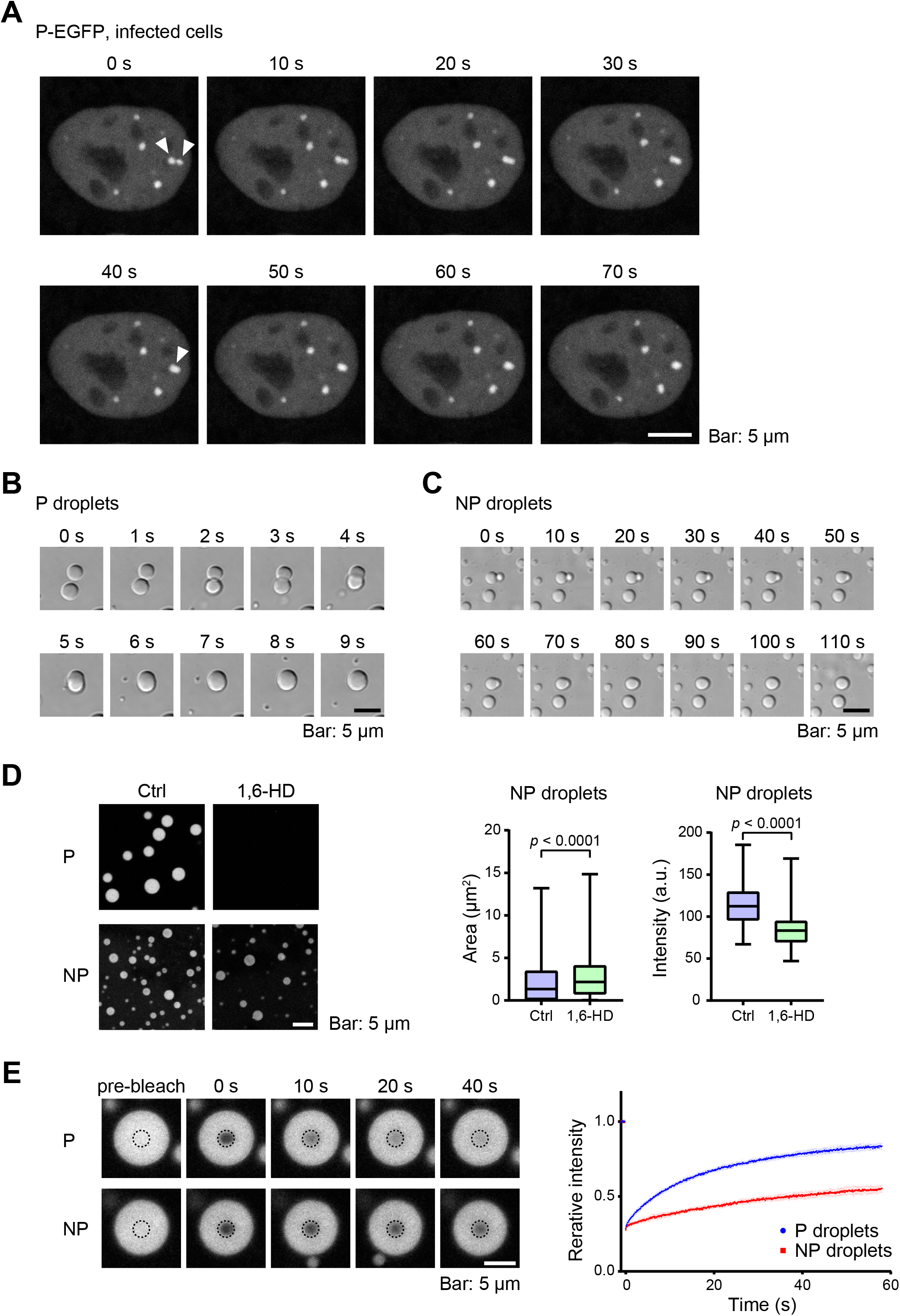
The liquid-like properties the BoDV-1 condensates. **A**. Time-lapse imaging of the fusion events of vSPOTs. P-EGFP was expressed in BoDV-1-infected OL cells and observed by confocal microscopy. **B** and **C**. Time-lapse DIC imaging of the fusion events of droplets formed by BoDV-1 P alone (**B**) or N and P (**C**). **D**. Dissolution of P and NP droplets by the addition of 6% 1,6-HD. The area and intensity of fluorescence per unit area were calculated. Statistical significance was calculated with the two-tailed unpaired *t*-test. **E**. Representative images in the FRAP experiment of P and NP droplets (left) and the recovery curves of the fluorescence intensity of P-EGFP after photobleaching (right). The intensities are presented as relative values for which the mean intensities before photobleaching were set to a value of 1.

Next, we investigated and compared the properties of the liquid droplets formed by P alone (P droplets) and by both N and P (NP droplets) by a simplified *in vitro* system. Fusion events were observed with the P and NP droplets (Fig. 2B and C), suggesting that both have liquid properties. However, the NP droplets required a much longer time for two droplets to form one spherical droplet as compared to the P droplets (Fig. 2B and C). Additionally, the P droplets were completely dissolved by 1,6-hexanediol (1,6-HD), which is an aliphatic alcohol that affects the formation of liquid droplets ^26^, while the NP droplets were relatively more resistant to 1,6-HD, although the size and solubility were reduced (Fig. 2D). FRAP analysis showed that the fluorescence recovery of P-EGFP was slower within the NP droplets than the P droplets (Fig. 2E). These results suggest that N decreases the fluidity and increases the stability of droplets.

### Both N and P are required for the formation of the condensates in cells

The BoDV-1 nuclear IBs (i.e., vSPOTs) contain several viral proteins (N, P, X, M, and L), viral RNA, and the host protein HMGB1 ^16,17^. Although other mononegaviruses, such as rabies virus, measles virus, RSV, metapneumovirus, and human parainfluenza virus 3, require the expression of both N and P for the formation of viral IBs in uninfected cells, the proteins needed for the formation of vSPOTs in cells remain unknown. Hence, we expressed N alone, P alone, and both N and P in uninfected U-2 OS cells and observed them by immunofluorescence staining and confocal microscopy. When N alone was expressed, condensates were not observed in the nucleus, although aggregates were observed in the cytoplasm (Fig. 3A). The expression of P alone did not induce the formation of nuclear condensates (Fig. 3B). On the other hand, the expression of both N and P induced the formation of nuclear condensates that were morphologically similar to vSPOTs (Fig. 3C, arrowheads). The nuclear condensates induced by the expression of N and P contained HMGB1, a host factor that is known to be localized to the vSPOTs of infected cells (Fig. 3C). Note that the condensates containing both N and P were observed in the nucleus as well as the cytoplasm (Fig. 3C, arrows). The expression of N and P’ did not induce the formation of condensates (Fig. 3D), which is consistent with the *in vitro* result (Fig. 1E). These results suggest that N and P are sufficient for the formation of BoDV condensates in cells, as also reported for other mononegaviruses, and P, but not P’, contributes to the formation of the condensates.

**Fig. 3.**
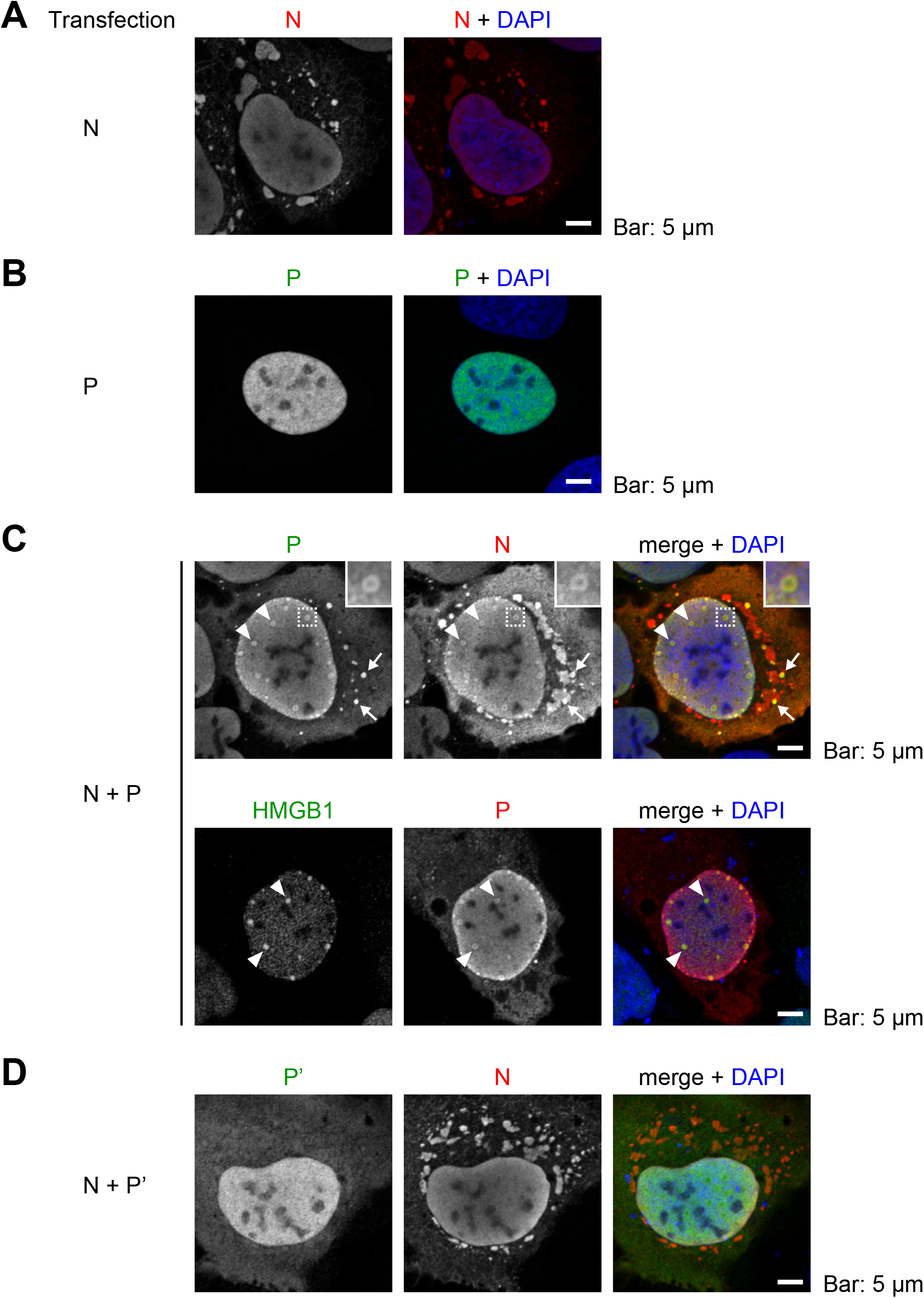
Condensates formed by BoDV-1 N and P in cells. BoDV-1 N alone (**A**), P (p24) alone (**B**), N and P (p24) (**C**), and N and P (p16) (**D**) were expressed in U-2 OS cells, which were subjected to immunofluorescence staining followed by confocal microscopy. The magnified images of the areas indicated by the rectangles are inserted (**C**, top).

### RNA controls the LLPS of BoDV-1 condensates

The N of mononegaviruses encapsidates RNA and forms nucleocapsids. P, also a component of RNPs, functions as a cofactor of L, which is a RNA-dependent RNA polymerase ^18,27^. From this perspective, there is a possible, although elusive, relationship between P and RNA. Therefore, we investigated possible interactions between P and the BoDV-1 minigenome RNA (mgRNA), which contains the leader and trailer of BoDV-1 and Gluc sequences (Fig. S3A) ^19^, with the use of the electrophoretic mobility shift assay. A band shift was observed with an increasing concentration of P, suggesting interactions with RNA (Fig. 4A). On the other hand, P’ did not interact with RNA. Note that a band shift was also observed when using RNA lacking the 3’ leader and 5’ trailer regions (Fig. S3B), suggesting that P interacts with RNA in a sequence-independent manner. No apparent shift was observed with the P proteins of other mononegaviruses (i.e., VSV and RSV) (Fig. 4A).

**Fig. 4.**
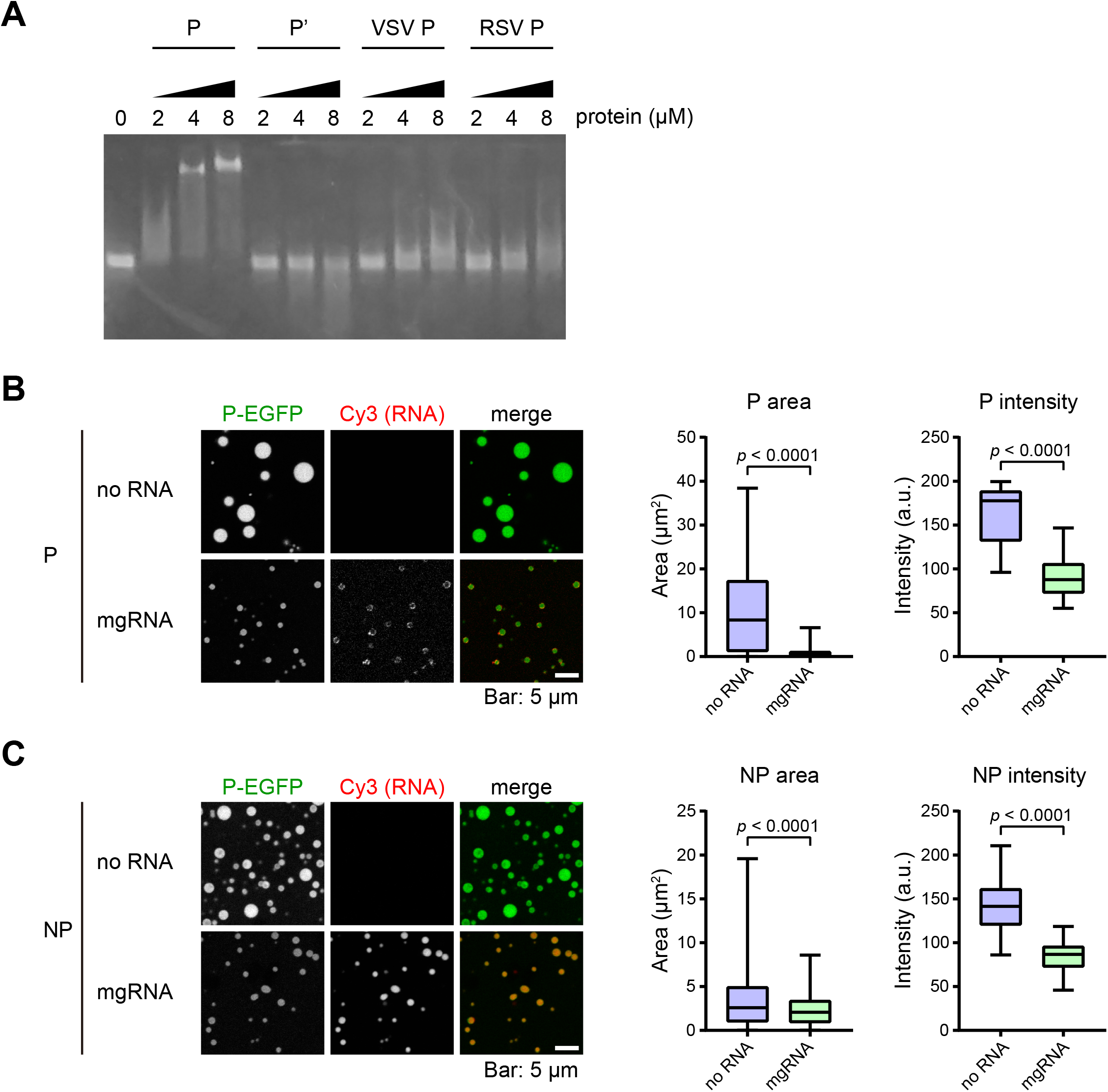
The regulation of the BoDV-1 liquid droplets by RNA. **A**. The indicated concentration of BoDV-1 P (p24), P (p16), VSV P, or RSV P was mixed with the BoDV-1 minigenome RNA and subjected to the electrophoretic mobility shift assay. **B** and **C**. Fluorescence signals of P (**B**) and NP (**C**) droplets were observed by confocal microscopy in the presence or absence of 200 ng/µl of Cy3-labeled BoDV-1 minigenome RNA. The area and intensity of fluorescence per unit area were calculated. Statistical significance was calculated with the two-tailed unpaired *t*-test.

We also investigated the possible incorporation of mgRNA into the droplets. We observed that mgRNA was incorporated into both the P and NP droplets (Fig. 4B and C), consistent with the RNA-binding activity of BoDV-1 P (Fig. 4A). However, the signal intensity of RNA was much stronger in the NP than P droplets, which probably reflects the role of BoDV-1 N in encapsidating viral genomic RNA.

RNA regulates the formation of liquid droplets formed by RNA-binding proteins ^28–31^. Therefore, we analyzed the effects of adding RNA to the P and NP droplets. The results showed that mgRNA inhibited the formation of not only P but also NP droplets (Fig. 4B and C, S1C). These results suggest that RNA is incorporated into the viral condensates in cells, but an excess amount of RNA could begin to dissolve the viral condensates (see Discussion).

## Discussion

Many mononegaviruses form IBs in infected cells. These IBs are thought to be the sites of viral transcription and replication, thus it is important to elucidate the properties and organizations of viral IBs to gain a deeper understanding of the replication strategies of viruses. Additionally, because viral IBs are biomolecular condensates formed by exogenous materials in cells, it is helpful to understand the properties of biomolecular condensates of organisms. In this study, to reveal the organization of condensates formed by BoDV-1, we analyzed the basic properties of BoDV-1 N and P. Similar to other cytoplasmic mononegaviruses, co-expression of BoDV-1 N and P was sufficient to reconstruct the viral IB-like structures in cells (Fig. 3). On the other hand, BoDV-1 P alone drove the formation of liquid droplets *in vitro*, which was dependent on the IDR (Fig. 1). This feature is distinct from other mononegaviruses that needs both N and P to form liquid droplets. Further, P of BoDV-1, but not other mononegaviruses (i.e., VSV and RSV), binds to RNA and high concentrations of RNA dissolved droplets formed by BoDV-1 N and P *in vitro* (Fig. 4 and S3). These findings revealed that although BoDV-1 shares common strategies with other mononegaviruses for the formation of IBs, at least to a certain extent, BoDV-1 apparently adopted strategies different from those of other mononegaviruses.

The unique feature of BoDV-1 P to form liquid droplets without N may be attributed to the overlap of the region important for driving LLPS with the RNA-binding region. We revealed that the N-terminal IDR is important for the RNA-binding activity of BoDV-1 P (Fig. 4), which is overlapped with the region necessary to drive LLPS *in vitro* (Fig. 1). Further, an excess amount of RNA dissolved the BoDV-1 P droplets (Fig. 4). These findings suggest that the LLPS and RNA-binding activities of BoDV-1 might be competing against each other. Notably, a high concentration of RNA is reportedly present in the cell nucleus ^28^. Thus, BoDV-1 P may have a greater ability to drive LLPS than the other P proteins of mononegaviruses to induce LLPS in the nuclear environment.

The transcription and replication of BoDV-1 might be regulated by the competitive feature of BoDV-1 P. We revealed that an excess amount of RNA also dissolved the NP droplets, even though RNA was efficiently incorporated into the NP droplet, which was probably due to the ability of BoDV-1 N to encapsidate RNA (Fig. 4). Considering that BoDV-1 may control transcription and replication to establish persistent infection in the nucleus, we propose the following hypothetical mechanism underlying the regulation of viral transcription and replication: if viral transcription and replication are excessive, the excess RNA binds to the N-terminal IDR of BoDV-1 P, leading to dissolving the viral IBs to arrest them. Although the detailed relationship between the LLPS driving and RNA-binding abilities of BoDV-1 P remain unclear, this hypothesis could explain one of the mechanisms needed to establish a persistent infection in the host cell.

Our data suggest that BoDV-1 N plays an important role in the formation of viral IBs in cells. Although our data suggest that BoDV-1 P has greater ability to drive LLPS, co-expression of BoDV-1 N was needed to induce the formation of viral IB-like structures in cells (Fig. 3). Importantly, we could compare the properties of the P and NP droplets *in vitro* because BoDV-1 P alone can form liquid droplets, which makes it possible to speculate the effect of BoDV-1 N on viral condensates. Notably, the NP droplets had lower fluidity and were more resistant to treatment with 1,6-HD than the P droplets (Fig. 2). Our previous studies revealed that BoDV-1 IBs had insoluble properties and that BoDV-1 N has an immobile property ^16,32^. Thus, besides the function as a nucleocapsid protein (i.e., encapsidation of viral genomic RNA), BoDV-1 N might also convey the properties of insolubility and/or viscosity to viral IBs, which might lead to stable transcription and replication.

In this study, we did not test the effects of viral proteins other than BoDV-1 N and P. As described above, other viral proteins, including X, M, L, are colocalized in the vSPOTs. Although our data indicated that both N and P are sufficient for the formation of condensates as phase-separated structures, other viral components may affect the formation, structure, localization, and function of viral condensates as reported for other mononegaviruses ^33^. Indeed, some of the condensates formed by BoDV-1 N and P were observed in the cytoplasm (Fig. 3), although cytoplasmic viral IBs were barely observed in BoDV-1-infected cells. Hence, further analyses are necessary to elucidate the organization and function of viral IBs.

LLPS is adopted not only in mononegaviruses but also in many other RNA viruses, including severe acute respiratory syndrome coronavirus 2 (SARS-CoV-2). In SARS-CoV-2, the nucleocapsid protein forms condensates by LLPS in an RNA-dependent manner ^30,31,34–38^. Remarkably, a higher amount of RNA was reported to dissolve the condensates formed by the nucleocapsid protein and RNA ^30,31,36,37^, which is consistent with the results of the present study (Fig. 4), as well as other cellular proteins ^28^. This suggests that regulation of the phase-separated condensates by RNA is a common mechanism not only among both positive-strand and negative-strand RNA viruses but also among cells. Further analyses focusing on LLPS are warranted to elucidate the various phenomena in the life cycle of RNA viruses, leading to the development of anti-viral reagents targeting phase-separated condensates.

Taken together, the results of this study will help to elucidate the mechanisms underlying the formation of phase-separated condensates, which should be useful to further develop antivirals and understand the cellular biology.

## Acknowledgments

We are grateful to Takehiro Kanda and Chiaki Tanaka (Kyoto University) for technical support. We would like to thank Enago (www.enago.jp) for the English language reveiw.

## Author contributions

YH and MH conceived and designed the study. YH and MH performed experiments. YH and MH analyzed the data, and all authors discussed the data interpretation. YH wrote the initial draft of the manuscript, and all authors revised the manuscript.

## References

1. Kuhn, J. H. et al.. 2020 taxonomic update for phylum Negarnaviricota (Riboviria: Orthornavirae), including the large orders Bunyavirales and Mononegavirales. Archives of Virology 165, (2020).

2. Nevers, Q., Albertini, A. A., Lagaudrière-Gesbert, C. & Gaudin, Y. Negri bodies and other virus membrane-less replication compartments. Biochim. Biophys. Acta - Mol. Cell Res. 1867, 118831 (2020).

3. Brangwynne, C. P. et al. Germline P Granules Are Liquid Droplets That Localize by Controlled Dissolution/Condensation. Science (80-.). 324, 1729–1732 (2009).

4. Molliex, A. et al. Phase Separation by Low Complexity Domains Promotes Stress Granule Assembly and Drives Pathological Fibrillization. Cell 163, 123– 133 (2015).

5. Berry, J., Weber, S. C., Vaidya, N., Haataja, M. & Brangwynne, C. P. RNA transcription modulates phase transition-driven nuclear body assembly. Proc. Natl. Acad. Sci. 112, E5237–E5245 (2015).

6. Altmeyer, M. et al. Liquid demixing of intrinsically disordered proteins is seeded by poly(ADP-ribose). Nat. Commun. 6, (2015).

7. Nikolic, J. et al. Negri bodies are viral factories with properties of liquid organelles. Nat. Commun. 8, 1–12 (2017).

8. Heinrich, B. S., Maliga, Z., Stein, D. A., Hyman, A. A. & Whelan, S. P. J. Phase transitions drive the formation of vesicular stomatitis virus replication compartments. MBio 9, 1–10 (2018).

9. Zhou, Y., Su, J. M., Samuel, C. E. & Ma, D. Measles Virus Forms Inclusion Bodies with Properties of Liquid Organelles. J. Virol. 93, 1–18 (2019).

10. Guseva, S. et al. Measles virus nucleo- and phosphoproteins form liquid-like phase-separated compartments that promote nucleocapsid assembly. Sci. Adv. 6, eaaz7095 (2020).

11. Galloux, M. et al. Minimal Elements Required for the Formation of Respiratory Syncytial Virus Cytoplasmic Inclusion Bodies In Vivo and In Vitro. MBio 11, 1–16 (2020).

12. Derdowski, A. et al. Human metapneumovirus nucleoprotein and phosphoprotein interact and provide the minimal requirements for inclusion body formation. J. Gen. Virol. 89, 2698–2708 (2008).

13. Zhang, S. et al. An amino acid of human parainfluenza virus type 3 nucleoprotein is critical for template function and cytoplasmic inclusion body formation. J Virol 87, 12457–12470 (2013).

14. Kuhn, J. H. et al. Taxonomic reorganization of the family Bornaviridae. Arch. Virol. 160, 621–632 (2015).

15. Cubitt, B. & de la Torre, J. C. Borna disease virus (BDV), a nonsegmented RNA virus, replicates in the nuclei of infected cells where infectious BDV ribonucleoproteins are present. J. Virol. 68, 1371–1381 (1994).

16. Matsumoto, Y. et al. Bornavirus closely associates and segregates with host chromosomes to ensure persistent intranuclear infection. Cell Host Microbe 11, 492–503 (2012).

17. Hirai, Y. et al. Borna disease virus assembles porous cage-like viral factories in the nucleus. J. Biol. Chem. 291, 25789–25798 (2016).

18. Schneider, U., Naegele, M., Staeheli, P. & Schwemmle, M. Active borna disease virus polymerase complex requires a distinct nucleoprotein-to-phosphoprotein ratio but no viral X protein. J. Virol. 77, 11781–9 (2003).

19. Reuter, A. et al. Synergistic antiviral activity of ribavirin and interferon-α against parrot bornaviruses in avian cells. J. Gen. Virol. 97, 2096–2103 (2016).

20. Nakamura, Y. et al. Isolation of Borna disease virus from human brain tissue. J. Virol. 74, 4601–11 (2000).

21. Kobayashi, T. et al. Modulation of Borna disease virus phosphoprotein nuclear localization by the viral protein X encoded in the overlapping open reading frame. J. Virol. 77, 8099–107 (2003).

22. Lin, Y., Protter, D. S. W., Rosen, M. K. & Parker, R. Formation and Maturation of Phase-Separated Liquid Droplets by RNA-Binding Proteins. Mol. Cell 60, 208–219 (2015).

23. Nott, T. J. et al. Phase Transition of a Disordered Nuage Protein Generates Environmentally Responsive Membraneless Organelles. Mol. Cell 57, 936–947 (2015).

24. Kobayashi, T., Watanabe, M., Kamitani, W., Tomonaga, K. & Ikuta, K. Translation initiation of a bicistronic mRNA of Borna disease virus: A 16-kDa phosphoprotein is initiated at an internal start codon. Virology 277, 296–305 (2000).

25. Charlier, C. M. et al. Analysis of borna disease virus trafficking in live infected cells by using a virus encoding a tetracysteine-tagged p protein. J. Virol. 87, 12339–48 (2013).

26. Lin, Y. et al. Toxic PR Poly-Dipeptides Encoded by the C9orf72 Repeat Expansion Target LC Domain Polymers. Cell 167, 789-802.e12 (2016).

27. Perez, M., Sanchez, A., Cubitt, B., Rosario, D. & de la Torre, J. C. A reverse genetics system for Borna disease virus. J. Gen. Virol. 84, 3099–104 (2003).

28. Maharana, S. et al. RNA buffers the phase separation behavior of prion-like RNA binding proteins. Science 360, 918–921 (2018).

29. Zhang, H. et al. RNA Controls PolyQ Protein Phase Transitions. Mol. Cell 60, 220–230 (2015).

30. Lu, S. et al. The SARS-CoV-2 nucleocapsid phosphoprotein forms mutually exclusive condensates with RNA and the membrane-associated M protein. Nat. Commun. 12, 502 (2021).

31. Iserman, C. et al. Genomic RNA Elements Drive Phase Separation of the SARS-CoV-2 Nucleocapsid. Mol. Cell 80, 1078-1091.e6 (2020).

32. Hirai, Y., Honda, T., Makino, A., Watanabe, Y. & Tomonaga, K. X-linked RNA-binding motif protein (RBMX) is required for the maintenance of Borna disease virus nuclear viral factories. J. Gen. Virol. 96, 3198–203 (2015).

33. Ringel, M. et al. Nipah virus induces two inclusion body populations: Identification of novel inclusions at the plasma membrane. PLoS Pathog. 15, (2019).

34. Perdikari, T. M. et al. SARS CoV 2 nucleocapsid protein phase separates with RNA and with human hnRNPs. EMBO J. 39, 1–15 (2020).

35. Savastano, A., Ibáñez de Opakua, A., Rankovic, M. & Zweckstetter, M. Nucleocapsid protein of SARS-CoV-2 phase separates into RNA-rich polymerase-containing condensates. Nat. Commun. 11, 6041 (2020).

36. Carlson, C. R. et al. Phosphoregulation of Phase Separation by the SARS-CoV-2 N Protein Suggests a Biophysical Basis for its Dual Functions. Mol. Cell 80, 1092-1103.e4 (2020).

37. Cubuk, J. et al. The SARS-CoV-2 nucleocapsid protein is dynamic, disordered, and phase separates with RNA. Nat. Commun. 12, 1936 (2021).

38. Zhao, M. et al. GCG inhibits SARS-CoV-2 replication by disrupting the liquid phase condensation of its nucleocapsid protein. Nat. Commun. 12, 2114 (2021).

39. Mészáros, B., Erdös, G. & Dosztányi, Z. IUPred2A: Context-dependent prediction of protein disorder as a function of redox state and protein binding. Nucleic Acids Res. 46, W329–W337 (2018).

