## Supplementary Figures for "The mechanism underlying the organization of Borna disease virus inclusion bodies is unique among mononegaviruses"

### Fig.S1

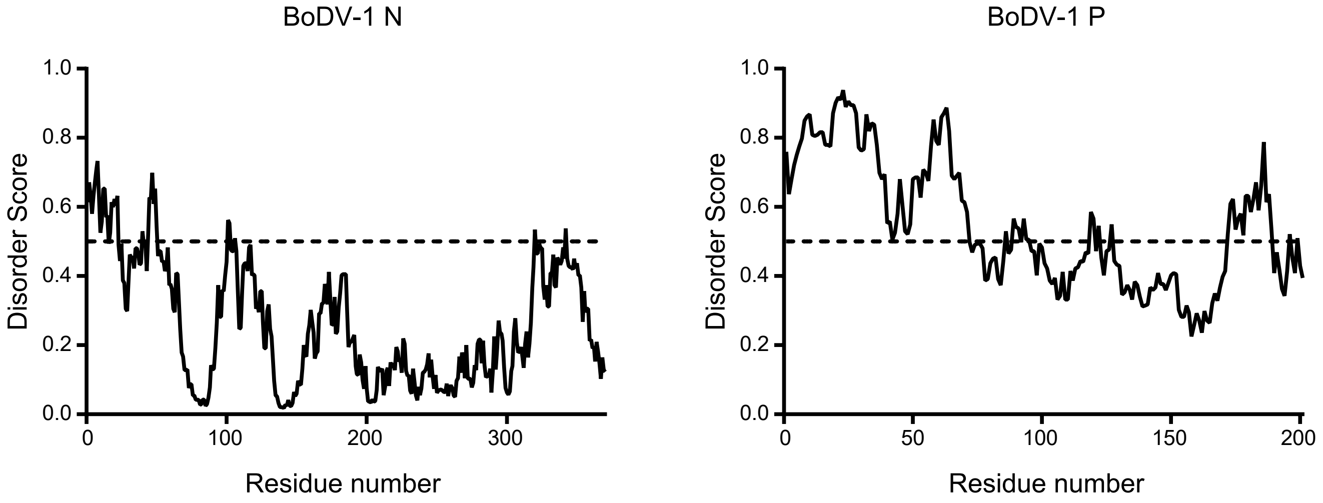


**Fig. S1 The prediction of intrinsically disordered regions (IDRs) of BoDV-1 N and P.**
The IDRs of BoDV-1 N and P were predicted in IUPred2A (Mészáros et al, 2018). The result for the prediction of P (right) is identical with Fig. 1A.

#### Fig.S2

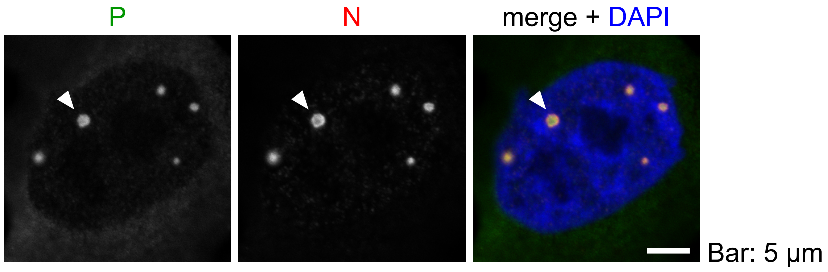


**Fig. S2 BoDV-1 forms inclusion bodies, termed as viral speckle of transcripts (vSPOTs) in the infected cells.**
BoDV-1-infected cells were subjected to immunofluorescence staining using anti-N and anti-P antibodies, followed by confocal microscopy. The arrowheads indicate the tipical vSPOTs.

### Fig.S3

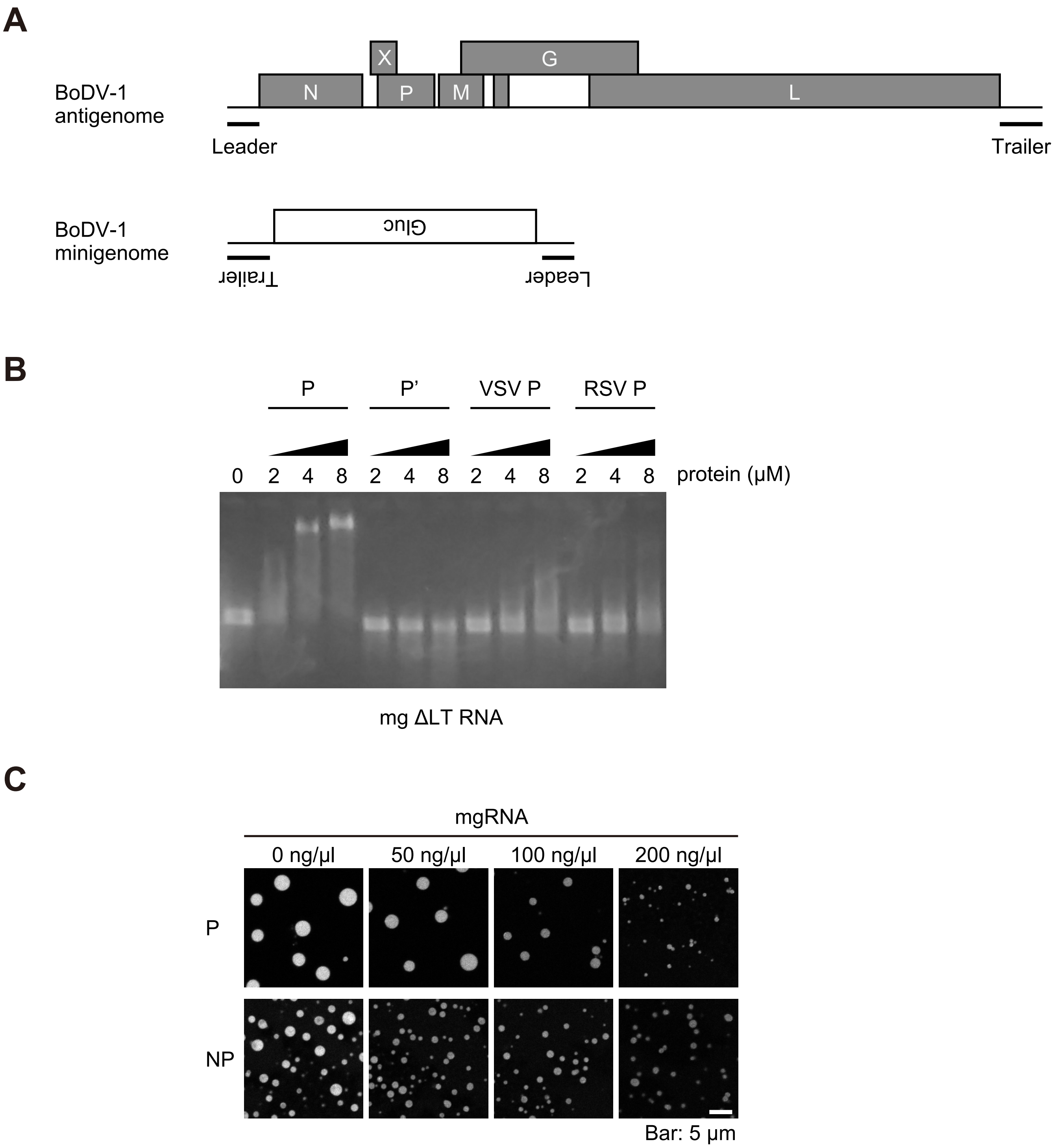


**Fig. S3 RNA-dependent regulation of the liquid droplets formed by BoDV-1 proteins.**
A. Schematic diagrams of BoDV-1 antigenome and minigenome. B. Indicated concentration of P, P' , VSV P and RSV P were mixed with mg RNA that lacks both 3' leader and 5' trailer regions (mg ΔLT RNA), and subjected to electrophoretic mobility shift assay (EMSA). C. Indicated concentration of Cy3-labeled mgRNA were added to the P and NP droplets that were fluorescently labeled by P-EGFP, and fluorescence signals were observed by confocal microscopy.
